# The thylakoid lumen Deg1 protease affects non-photochemical quenching via the levels of violaxanthin de-epoxidase and PsbS

**DOI:** 10.1101/2024.05.14.594122

**Authors:** Elinor Aviv-Sharon, Laure D. Sultan, Leah Naveh, Dana Charuvi, Meital Kupervaser, Ziv Reich, Zach Adam

## Abstract

Non-photochemical quenching (NPQ), the dissipation of excess light energy as heat, has been long recognized as a major protective mechanism that minimizes the potential for oxidative damage to photosystem II (PSII) reaction centers. Two major positive contributors to NPQ are the carotenoid zeaxanthin, generated from violaxanthin by the enzyme violaxanthin de-epoxidase (VDE or NPQ1), and the thylakoid protein PsbS (NPQ4). The involvement of the lumenal Deg proteases in the repair of PSII from photoinhibition prompted us to further explore their possible role in other responses of *Arabidopsis thaliana* to high light. Here we show that upon exposure to high light, the single *deg1* and the triple *deg158* mutants display different levels and kinetics of NPQ, compared to the *deg58* mutant and WT that behave alike. In response to high light, the two genotypes lacking Deg1 over-accumulate NPQ1 and NPQ4. After temporal inhibition of protein translation in vivo, the level of these two proteins in *deg1* is higher than in WT. Together, the results suggest that Deg1 represents a new level of regulation of the NPQ process through adjusting the quantity of NPQ1 and NPQ4 proteins, probably through their proteolysis.

## INTRODUCTION

Light energy absorbed by photosynthetic pigments and transferred to photosynthetic reaction centers (RCs) drives photosynthesis and hence life on Earth. Only a fraction of the absorbed light is utilized for photochemistry, while the rest is released mostly as heat, and also as fluorescence. Chlorophyll fluorescence has been traditionally used as a non-destructive tool for monitoring different parameters of photosynthesis (Maxwell and Johnson, 2000). Accordingly, photochemistry, the conversion of light energy to chemical one, occurring in RCs, has been referred to as ‘photochemical quenching’ of chlorophyll fluorescence, whereas the excess energy dissipated as heat was coined ‘non-photochemical quenching’ (NPQ) (Ruban et al., 2012; Ruban, 2016). Under natural conditions, the relative amount of energy driving photosynthetic electron transfer (ETR) and that released as heat vary. In fact, channeling excess excitation energy to NPQ has been considered a protective mechanism that minimizes the potential for oxidative damage to RCs, mostly to those of photosystem II (PSII) (Ruban et al., 2012; Nicol et al., 2019). Such damage, the phenomenon known as ‘photoinhibition’ (Powles, 1984; Adir et al., 2003), leads to decreased photosynthetic activity, which can affect plant growth and fitness.

NPQ is triggered upon acidification of the thylakoid lumen during photosynthetic ETR. Two major components contribute to NPQ in higher plants: the carotenoid zeaxanthin and the protein PsbS. The precise mechanism of how zeaxanthin mediates NPQ is not clear. However, its level is determined by the balance between the activity of two enzymes of the xanthophyll cycle: violaxanthin de-epoxidase (VDE, also known as NPQ1) that converts violaxanthin to zeaxanthin, and zeaxanthin epoxidase (ZEP or NPQ2), which mediates the opposite reaction (Niyogi et al., 1998; Jahns and Holzwarth, 2012). PsbS (also known as NPQ4) is a thylakoid membrane protein that does not bind pigments (Li et al., 2000; Li et al., 2002). Its activation involves light-induced monomerization, that is accompanied by reorganization of the PSII antenna and binding to LHCII trimers (Correa-Galvis et al., 2016).

Despite NPQ and other protective mechanisms, the PSII reaction center and especially its D1 protein are constantly prone to oxidative damage that leads to photoinhibition. Concomitant with this, a repair cycle that involves de-novo synthesis of D1 and other components of PSII operates in chloroplasts. Inherent to this cycle is the proteolytic degradation of oxidatively damaged proteins, which is carried out cooperatively by the thylakoid FtsH protease complex (Lindahl et al., 2000; Bailey et al., 2002; Zaltsman et al., 2005; Kato et al., 2012), facing the stromal side of the membrane, and the lumenal Deg1 and Deg5-Deg8 proteases (Kapri-Pardes et al., 2007; Sun et al., 2007; Kley et al., 2011). The apparently redundant function of Deg1 and Deg5-Deg8 was investigated through analysis of single, double and triple *deg* mutants, to uncover the physiological dominance of the former due to higher abundance and more potent proteolytic activity (Butenko et al., 2018). Thus, all mutants lacking Deg1 were smaller and more sensitive to stresses, whereas plants missing Deg5-Deg8 were very similar to WT plants under all tested conditions.

The involvement of the lumenal Deg proteases in the repair of PSII from photoinhibition prompted us to further explore their possible role in other responses of *Arabidopsis thaliana* to high light. In this study, we show that the single *deg1* and the triple *deg158* mutants exhibit different levels and kinetics of NPQ upon exposure to high light, compared to the *deg58* mutant and WT that behaved alike. Specifically, upon exposure to high light, they had higher levels of NPQ and slower recovery. Comparative proteomics revealed that the lack of either Deg1 or Deg5-Deg8 complexes was not compensated for by upregulation of the other. However, the two genotypes lacking Deg1 over-accumulated NPQ1 and NPQ4 upon exposure to high light. After temporal inhibition of protein translation in vivo, the level of these two proteins in *deg1* was higher than in WT. Taken together, the results suggest that Deg1 is involved in the regulation of the NPQ process through an effect on the level of NPQ1 and NPQ4 proteins.

## RESULTS

### Effect of high light on photosynthesis parameters in Deg protease mutants

The well-established role of Deg proteases in the response of Arabidopsis plants to photoinhibition (Kapri-Pardes et al., 2007; Sun et al., 2007; Kley et al., 2011; Kato et al., 2012) prompted us to explore possible additional roles of Deg in the response to high light (HL). To this end, we grew WT and Deg mutant plants, lacking either Deg1 (*deg1*), Deg5-Deg8 (*deg58*) proteases or both (*deg158*) in short days, under normal light (NL; ∼75 μmol photons m^-2^ s^-1^). Five-weeks old plants were transferred to HL (∼750 μmol photons m^-2^ s^-1^) for 1-8 h, or allowed to recover for additional 24 h in NL, and subjected to pulse-amplitude-modulation (PAM) measurements at each of these time points. The analyzed plants and their PAM images are provided in Supp. Fig. 1A. Consistent with our previous work (Butenko et al., 2018), prior to exposure to HL, *deg1* and *deg158* demonstrated a lower maximum efficiency of PSII (*Fv/Fm*) compared to WT and *deg58* (Supp. Fig. 1B). Upon exposure to HL, all four genotypes showed a continuous decrease in *Fv/Fm* values. However, the decrease in *deg1* and *deg158* was more pronounced, indicating that they were more sensitive to HL than WT and *deg58*. After 24 h of recovery under NL, all genotypes regained their original maximum efficiency of PSII. Consistent with this were the light response curves of photosynthetic electron transport rates (ETR) (Supp. Fig. 2), which also correlated to plant size of the four genotypes (Supp. Fig. 1A). Induction and relaxation curves of NPQ measured in the same plants provided some interesting insights. As shown in Fig. 1, NPQ levels in *deg1* and *deg158* plants were generally higher compared to WT and *deg58* plants. The rate of NPQ induction in the Deg1-lacking genotypes appeared to be faster, and the relaxation slower, than in the genotypes containing Deg1. Another interesting observation was the relation between the length of HL exposure and NPQ kinetics. In WT and *deg58*, the longer the exposure to HL was, the lower the NPQ levels were, up to 4 h of HL, suggesting an adaptation to the stress condition. Exposure to HL for 8 h did not result in further decrease in these two genotypes. This behavior is consistent with a previous report showing a lower contribution of NPQ under prolonged exposure to high light (Saccon et al., 2022). In contrast, no decrease in NPQ was observed in *deg1* and *deg158* plants. During 1-4 h of HL, the kinetics of NPQ remained the same. Only after exposure to 8 h of HL, the values of NPQ increased in plants lacking Deg1. After 24 h recovery under NL, the capacity for NPQ was restored in WT and *deg58*, although with somewhat different kinetics than before recovery. In *deg1* and *deg158* plants, the values of NPQ further increased after the recovery, to values well above those observed at the beginning of the experiment (Fig. 1). These results imply that Deg1 might somehow be involved in the regulation of NPQ.

**Figure 1.**
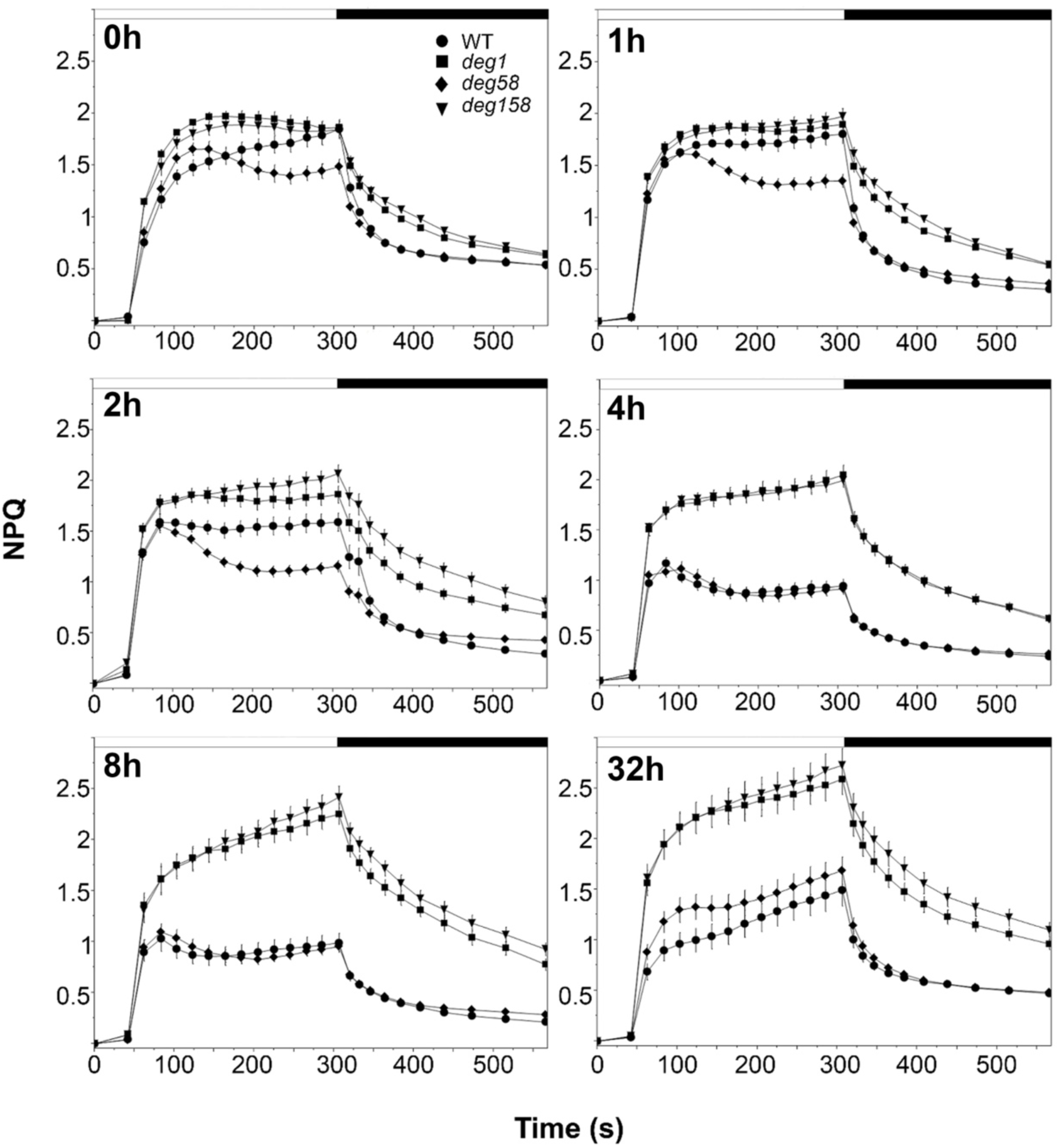
Induction/relaxation curves of non-photochemical quenching (NPQ). Recordings were performed on intact leaves before (0h) and following exposure to high-light (HL) (∼750 μmol m^2^ s^1^) for 1, 2, 4 and 8 hours (2h, 4h and 8h, respectively). (32h) Curves recorded on leaves exposed to HL for 8 h and then let to recover for 24 h under NL conditions. Values shown represent the mean ±SE obtained from recordings of 25 leaves from at least 3 different plants, at each time point, for each genotype.

### Effect of HL on the proteome of WT and Deg protease mutants

To further explore the effect of HL on WT and Deg protease mutants, we carried out a comparative proteomic analysis on the same set of four genotypes, at six different time points along the exposure to HL and following recovery at NL, with four biological replicates each. Mass-spectrometry (MS) analysis of the 96 samples allowed identification and quantification of 40,546 peptides, mapped to 4,188 different proteins (Supp. Table 1). Of these, ∼1,400 were predicted to be chloroplast proteins. Assuming that detectability is proportional to protein abundance, and that chloroplast proteins account for 10-15% of the plant proteome, these numbers indicate that chloroplast proteins are more abundant than those located in other compartments of the leaf cells. More than half of the chloroplast proteins were identified, compared to only ∼1/5 of the proteins residing in other compartments. Volcano plots of the data obtained from plants prior to exposure to HL revealed the extent of alterations in protein abundance only due to knocking out the Deg proteases. Cutoffs for significance were 2-fold change in the level of a given protein and p-value <0.05 in comparisons between the mutants and WT. As shown in Fig. 2, all three mutant genotypes demonstrated alterations in accumulation of proteins when compared to the WT proteome. Interestingly, changes were not limited to chloroplast proteins, as the numbers of over- or under-accumulating proteins among ‘all proteins’ (upper panels) were higher than those considered ‘chloroplast proteins’ (lower panels), suggesting that variations in the chloroplast proteome were transduced to impact other cellular compartments as well. Another interesting observation was that the levels of neither Deg1 nor Deg5-Deg8 were adjusted in the absence of the other Deg proteins, implying that the loss of one was not compensated for by over-accumulation of the other (Fig. 2). Moreover, pairwise Pearson’s correlation analysis on ‘time 0’ samples (NL, prior to exposure to HL) revealed positive high correlation between the proteomes of *deg1* and *deg158* plants, and lower but still positive correlation between WT and *deg58* plants (Supp. Fig. 3).

**Figure 2.**
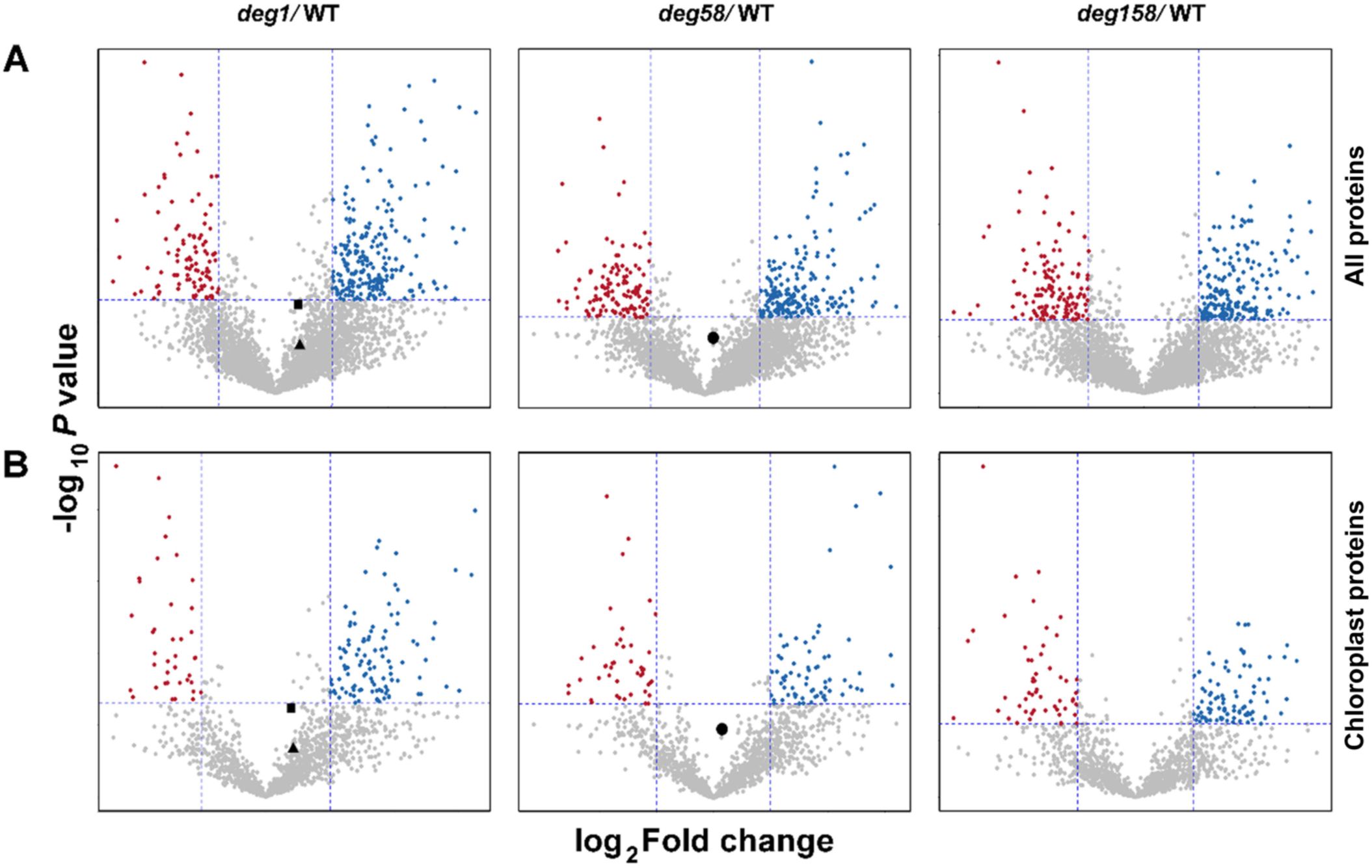
Volcano plots of proteomic data obtained from plants under normal light (NL) conditions. Volcano plots are depicted with the log_2_ fold-change (−1, 1; vertical dashed lines) of each protein and the -log_10_ p-value (p<0.05) (horizontal dashed line). The averages of the proteomic expression data of each mutant (*deg1*, *deg58* N = 4; *deg158*, N = 3) were compared with the averages of the data for WT (N = 4). Blue circles mark proteins whose level is higher in the mutants, whereas red circles mark those whose level is lower. Gray circles mark proteins whose level is not significantly altered or those which exhibit a log_2_ fold-change < 1. Deg1 (black circle), Deg5 (black triangle) and Deg8 (black square) are highlighted according to their p-values and fold change. Data are shown for (A) all 4188 identified proteins and (B) ∼1400 chloroplast proteins.

The relatedness of WT and *deg58* proteomes and those of *deg1* and *deg158* was retained also after exposure to HL. As can be seen by the principle component analysis (PCA) presented in Supp. Fig. 4, the different samples were clearly differentiated from each other at all time points, and the aforementioned pairs were grouped separately along PC1.

### The response of thylakoid proteins to HL in WT and Deg protease mutants

As Deg1 and Deg5-Deg8 proteases are found soluble in the thylakoid lumen, or peripherally attached to the lumenal-side of the thylakoid membrane, they can potentially degrade or cleave lumenal proteins or lumen-exposed regions of thylakoid membrane integral proteins. We therefore focused on the fate of such proteins along the exposure of WT plants to HL. Of the ∼1,400 identified proteins predicted to localize to chloroplasts, 118 were categorized as thylakoid integral membrane proteins, thylakoid peripheral proteins associated with the lumen side of the membrane, or soluble lumen proteins (Supp. Table 1, analyzed proteins – column ‘GI’). Fifty-five of these proteins were found differentially accumulating (FC> ±2, P value <0.05) in response to HL in WT. Nineteen proteins were up-regulated, including allene oxide synthase, FIB1a and FIB1b, all three involved in jasmonic acid (JA) biosynthesis, two components of the Cyt *b_6_f* complex, six proteins of the PSI core complex and its antenna, HCF244 involved in translation initiation of PsbA, PPD6 involved in redox regulation, CURT1A that controls grana architecture, and RIQ1 and RIQ2 involved in LHCII organization, as well as a number of proteins with unknown function (Supp. Table 1). The numerous down-regulated proteins in WT included the following: two subunits of ATP synthase, seven subunits of PSII, two subunits of PSI, 15 proteins involved in the assembly, stability and repair of PSII, the kinase STN8, the TLP18.3 phosphatase, FtsH1,2,5 and 8 proteases, and finally, NPQ1 (VDE) – the xanthophyll cycle enzyme synthesizing zeaxanthin.

To elucidate the impact of the absence of Deg1 and Deg5-Deg8 on the response of thylakoid membrane and lumenal proteins to HL, we performed a comprehensive co-expression clustering analysis. The 118 proteins identified as thylakoid integral membrane or lumenal proteins were subjected to K-means clustering, using Pearson’s correlation as a distance metric, and were grouped into nine clusters of co-regulated proteins in a genotype- and time-dependent manner (Fig. 3, Supp. Table 1). As depicted in the upper dendrogram of Fig. 3, WT and *deg58* samples were clearly separated from *deg1* and *deg158* samples, indicating that the effect of genotype on the accumulation of these proteins was stronger than the effect of HL. Of particular interest were clusters 2 and 4, containing proteins that over-accumulated in *deg1* and *deg158* at all time points. As such, at least some of these proteins may be substrates of the Deg1 protease. Proteins in cluster 2 included three subunits of NADH dehydrogenase, four factors associated with PSI or PSII assembly, FtsH1 and FtsH5 proteases, tocopherol synthase (VTE1), and most interestingly, NPQ1 (VDE). Cluster 4 proteins included the three aforementioned proteins associated with JA biosynthesis, the light-induced serine protease SppA, and NPQ4 (PsbS).

**Figure 3.**
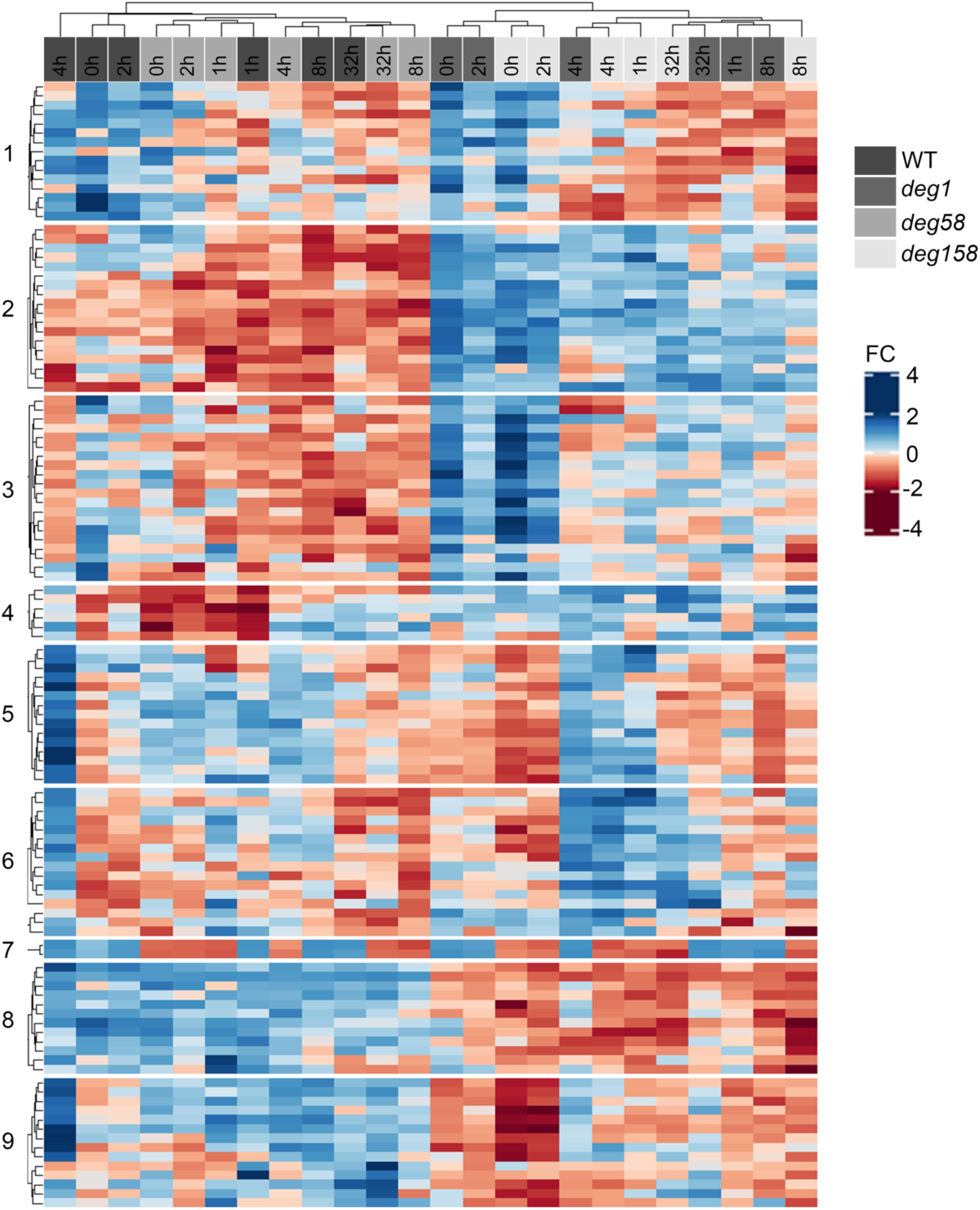
Differential expression of thylakoid membrane and lumenal proteins in WT and *deg* mutants under normal- and high light, and following recovery. The heat map generated for the thylakoid membrane and lumenal proteins identified in our study reveals segregation into nine co-expression groups.

Pairwise comparisons between the three mutant genotypes and WT plants, across the entire dataset, allowed us to focus on the 118 thylakoid proteins that might be in physical contact with the lumenal Deg proteases. As can be seen in Fig. 4, only very few thylakoid proteins over-accumulated in *deg58*. In contrast, 43 proteins were found to be up-regulated in many of the 12 remaining pairwise comparisons of *deg1* and *deg158* to WT, 22 of them in five comparisons or more. These included proteins involved in photosynthesis (plastocyanin, PPD3, PsbO2), PSII assembly/stability/repair (PSB33, HCF136, ALB3, FKBP-20-2, TLP18.3), NADPH dehydrogenase (PnsL1, PnsL4, PnsL5), plastoglobule proteins (FIB1b, FIB4), the protease SppA, uncharacterized proteins (TL20.3, AT1G32220, AT5G52970, HBP5, ENH1, STR10), and most noteworthy - NPQ1 and NPQ4 (Fig. 4). As enhanced expression of the genes encoding NPQ1 and NPQ4 could contribute to their over-accumulation, we determined their transcript levels at different time points during the exposure to HL. However, we did not detect differential massive increase in the level of these transcripts in *deg1* and *deg158* compared to WT (Supp. Fig. 5). Thus, the over-accumulation of these two proteins in the two genotypes lacking Deg1 during exposure to HL suggests that they might be substrates of this protease. The relatively higher levels of NPQ1 and NPQ4 may lead to the different level and kinetics of NPQ observed in *deg1* and *deg158* (Fig. 1).

**Figure 4.**
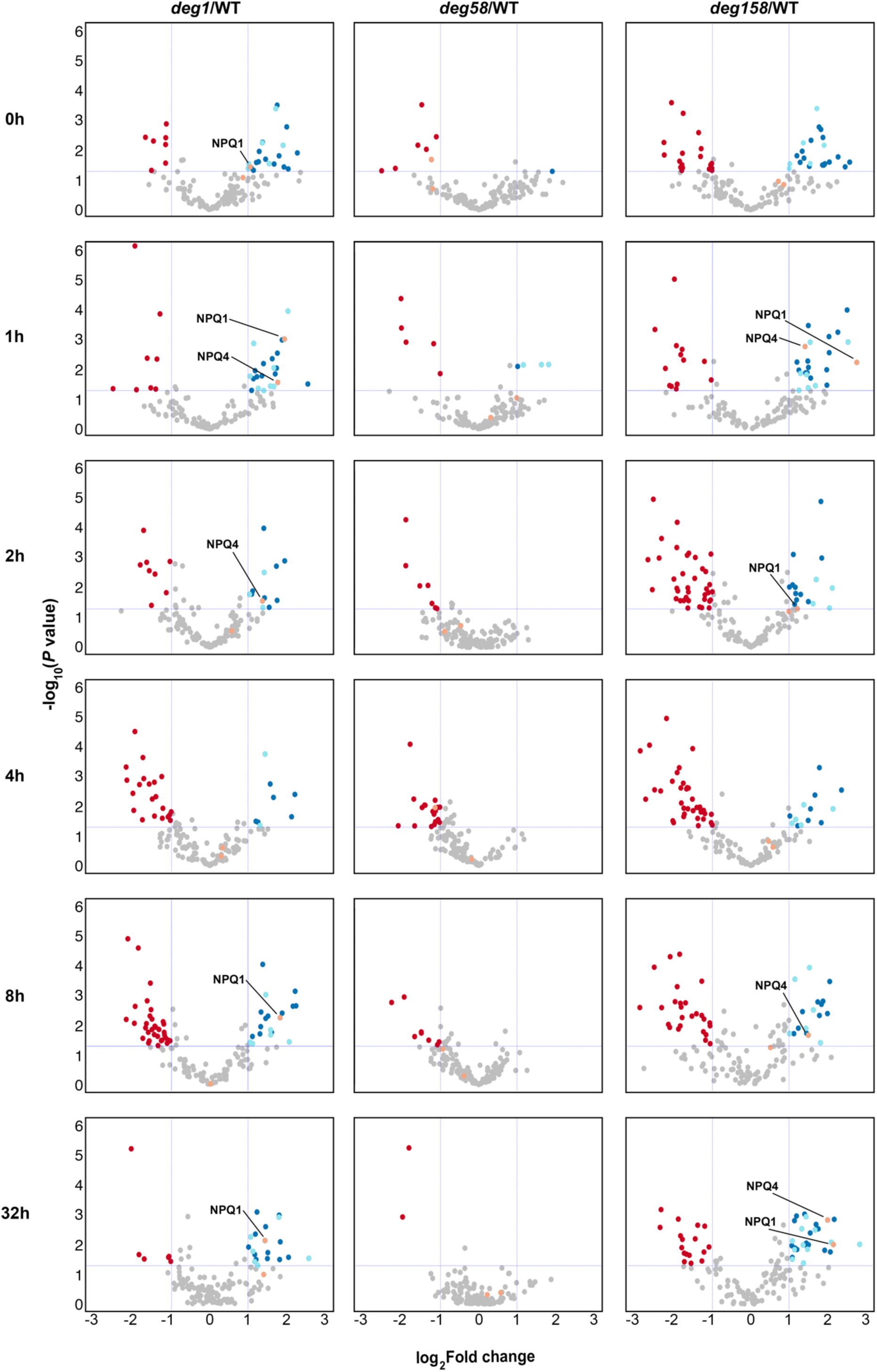
Volcano plots of thylakoid integral membrane and lumenal proteins expression levels in *deg* mutants *vs.* WT. Volcano plots are depicted with the log_2_ fold-change (vertical dashed lines) and the -log_10_ p-value (p<0.05) (horizontal dashed line) of the 118 proteins localized to the thylakoid membranes and lumen. The averages of the proteomic expression data of each mutant were compared with those of the WT at each time point. Blue and light blue circles mark lumenal and integral membrane proteins, respectively, whose levels increased more in the mutants relative to WT. Red circles denote proteins whose levels decreased more in the mutants vs. WT. Orange circles denote NPQ1 and NPQ4 proteins and highlight their over-accumulation in *deg1* and *deg158* plants compared to WT. Gray circles are proteins without any significant differences and/or log_2_ fold-change <1.

### The effect of cycloheximide on NPQ1 and NPQ4 in WT and the *deg1* mutant

To further explore the possibility that the level of NPQ1 and NPQ4 is regulated by Deg1, we tested the short-term effect of the protein translation inhibitor cycloheximide (CHX) on WT and *deg1* seedlings. To mitigate the size differences between WT and *deg1* plants observed in long-term autotrophic growth (see Supp. Fig. 1), seedlings of both genotypes were grown on Murashige & Skoog agar plates supplemented with 1% sucrose for 15 days. These seedlings, of almost similar size and appearance, were transferred to liquid medium, and incubated with or without CHX for 24 h. Before and after the incubation, total protein extracts were prepared from both genotypes and subjected to MS analysis. More than 5,000 different proteins were identified, 4,683 of them with at least two peptides, and these were quantified in the six different groups (two genotypes, before and after incubation with or without CHX; Supp. Table 2). As can be seen in the volcano plots (Fig. 5) and the heatmap (Supp. Fig. 6), the level of more than 700 proteins was altered in the WT in response to the CHX treatment. The level of 442 proteins decreased, whereas that of 268 proteins increased. Nevertheless, NPQ1 and NPQ4 were not among them. In the *deg1* mutant, over 1,000 proteins were affected by the presence of CHX. 475 proteins were down-regulated, and 578 proteins were up-regulated, among them were NPQ1 and NPQ4 (Fig. 5, Supp. Fig. 6 – cluster no. 3 and Supp. Table 2, analyzed proteins – column ‘BB’).

**Figure 5.**
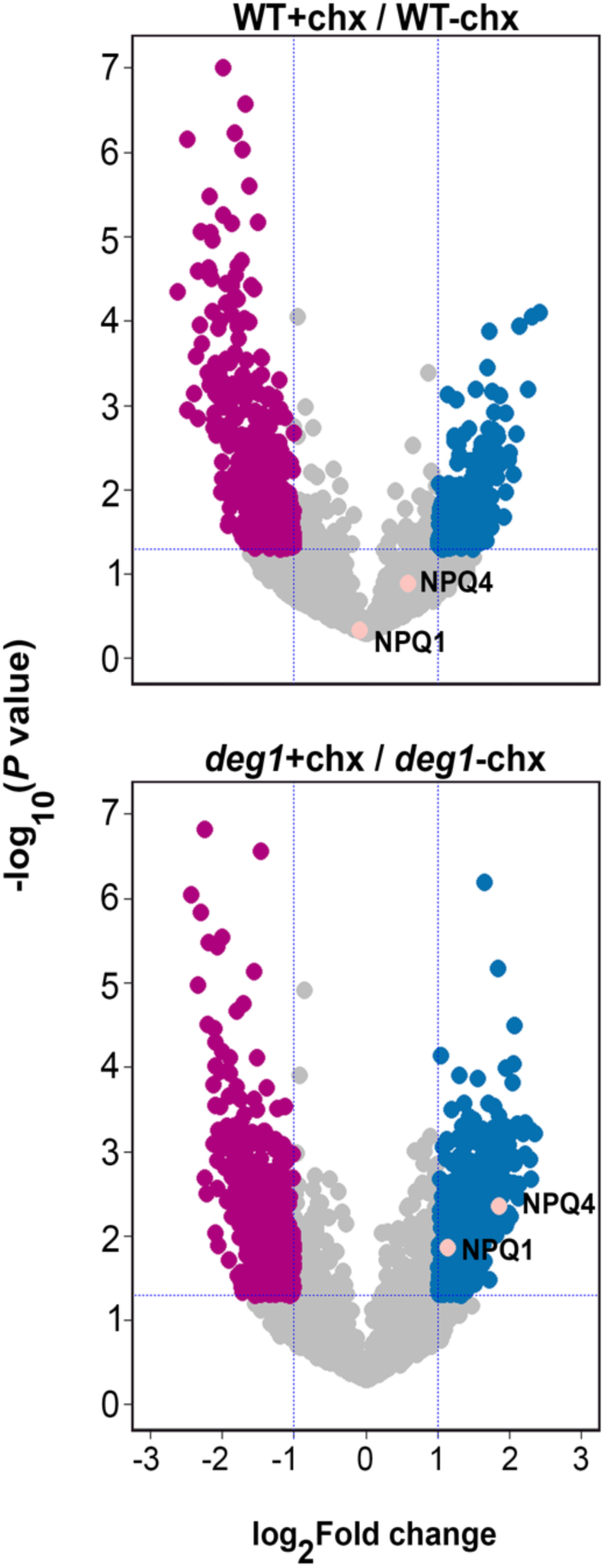
Volcano plot of proteomic data obtained from WT and *deg1* plants in response to cycloheximide treatment. Volcano plots are depicted with the log_2_ fold-change (vertical dashed lines) of each protein and the -log_10_ p-value (p<0.05) (horizontal dashed line). The averages of the proteomic expression data of *deg1* mutant and WT following 24-hour incubation with cycloheximide were compared with the averages of the data for *deg1* mutant and WT without cycloheximide incubation (N = 5). Blue circles mark proteins whose level was higher following the cycloheximide treatment, whereas magenta circles mark those whose level was lower. Gray circles mark proteins whose level was not significantly altered or those which exhibit a log_2_ fold-change < 1. NPQ1 and NPQ4 (orange circles) are highlighted according to their p-values and fold change.

Monitoring the level of the highly abundant stromal protein RBCL revealed no major differences in its level in the different treatments and genotypes (Fig. 6). Similarly, no differences in the level of NPQ1 and NPQ4 in the different samples were observed in the WT. However, after 24 h in the liquid medium, seedlings lacking Deg1 contained apparently more NPQ1, and statistically significant higher level of NPQ4 compared to WT. This trend was even more pronounced in the presence of CHX (Fig. 6). The similar, though somewhat less pronounced trend observed in the absence of CHX suggests that the over-accumulation of these proteins might be related to higher sensitivity of the *deg1* genotype to the transfer from solid to liquid culture, that might induce increased synthesis of the two proteins, before CHX penetrated the cells to exert its inhibitory effect. Nevertheless, the higher accumulation of both NPQ1 and NPQ4 in plants lacking Deg1 is consistent with the suggestion that the level of the two proteins is under the control of the Deg1 protease, most likely through their degradation by Deg1.

**Figure 6.**
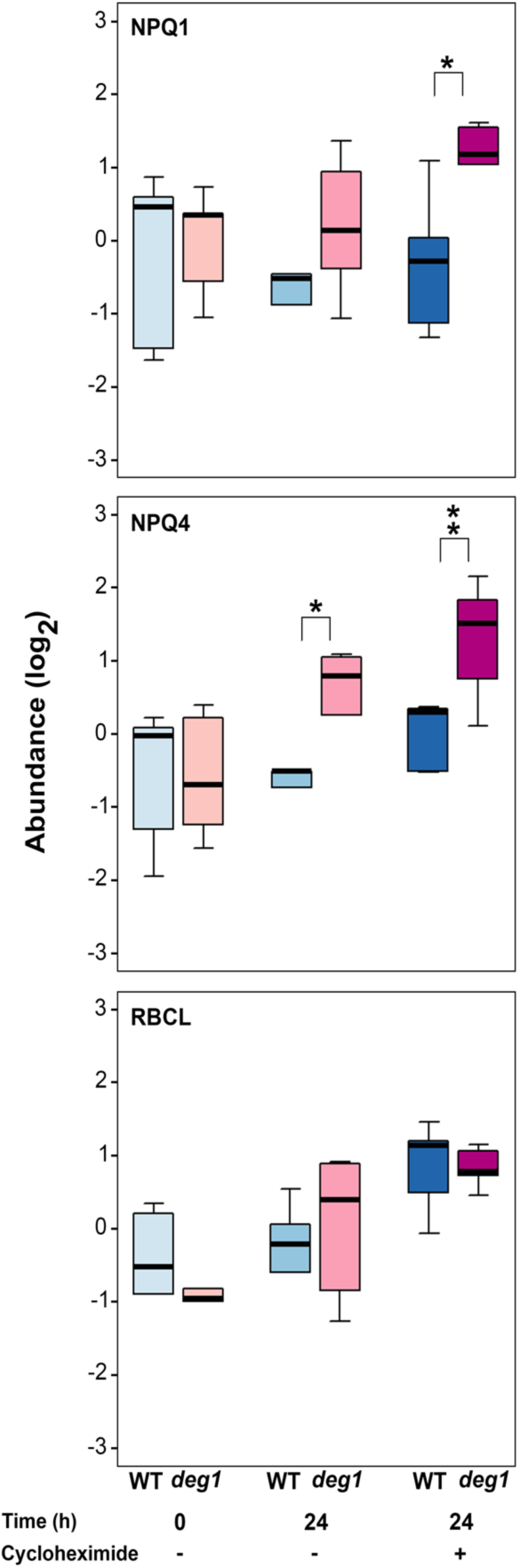
Effect of cycloheximide on accumulation of NPQ1 and NPQ4. Boxplots showing the levels of NPQ1, NPQ4 and RBCL proteins in WT and *deg1* mutant plants (N=5) before (0 h), with or without cycloheximide treatment for 24 h. *p<0.05, **p<0.01.

## DISCUSSION

NPQ has been long considered as a prominent mechanism protecting the photosynthetic machinery from oxidative damage through dissipation of excess light energy as heat. It is well established that the xanthophyll zeaxanthin, and the enzyme VDE (NPQ1) synthesizing it, as well as the PsbS (NPQ4) protein, are key components necessary for the induction of NPQ (Niyogi et al., 1998; Li et al., 2000; Li et al., 2002; Jahns and Holzwarth, 2012; Ruban et al., 2012; Ruban, 2016). Likewise, the enzyme ZEP (NPQ2), converting zeaxanthin to violaxanthin, and a number of other proteins, including SOQ1, LCNP and ROQH1, are involved in regulating the relaxation of NPQ (Brooks et al., 2013; Malnoe et al., 2018; Amstutz et al., 2020), enabling the funneling of more excitation energy to photochemistry upon the decline in light intensity. Expediting NPQ relaxation has been shown to increase plant biomass under fluctuating light conditions in tobacco (Kromdijk et al., 2016), although this was not the case for Arabidopsis (Garcia-Molina and Leister, 2020). The exact site where NPQ occurs is still a matter under investigation (Sacharz et al., 2017; Nicol et al., 2019).

The results of the current study suggest that the thylakoid lumen-located Deg1 protease is also involved in regulating the level of NPQ, through fine-tuning the levels of VDE and PsbS by proteolytic degradation. Three lines of evidence support this: i. The two genotypes that do not contain the Deg1 protease, *deg1* and *deg158*, accumulate higher levels of VDE (NPQ1) and PsbS (NPQ4) not only under optimal growth conditions (Butenko et al., 2018), but also upon exposure to HL (Figure 4); ii. These plants exhibit higher levels of NPQ compared to WT and plants lacking Deg5-Deg8 (Fig. 1). Consistent with this, ETR in the former genotypes is lower than in the latter (Supp. Fig. 2), and consequently, the mutant plants lacking Deg1 are smaller (Supp. Fig. 1; (Butenko et al., 2018)); iii. When protein synthesis is temporally inhibited, the level of NPQ proteins in *deg1* is higher than in WT (Fig. 6). This suggestion is also consistent with previous observations: recombinant Deg1 could bind PsbS (as well as a number of other thylakoid proteins from solubilized thylakoids), in a pull-down assay; and when recombinant Deg1 was incubated with solubilized thylakoids, PsbS was slightly degraded (Zienkiewicz et al., 2012). Thus, under natural conditions, when light intensity fluctuates, the balance between synthesis of VDE and PsbS and their proteolytic degradation determines how much excitation energy is funneled to photochemistry and how much is dissipated as heat (Fig. 7). In mutants lacking the Deg1 protease, the balance between synthesis and degradation of VDE and PsbS is impaired, leading to a higher level of heat dissipation and lower rates of ETR compared to WT, resulting in a decrease in photosynthesis yield (Fig. 7).

**Figure 7.**
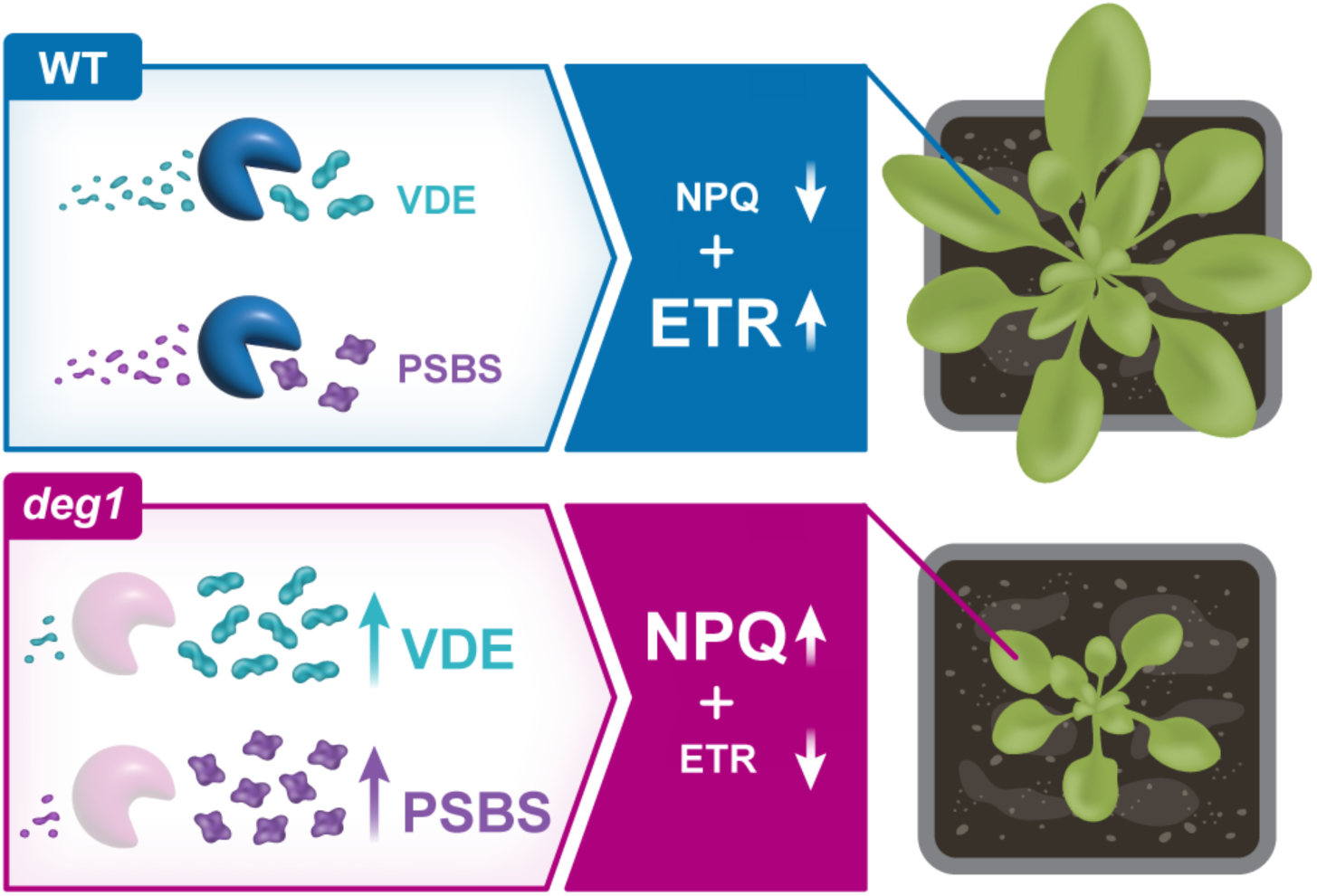
A model for the proposed role of the Deg1 protease in regulation of NPQ. In the presence of the thylakoid lumen-located Deg1 protease, NPQ levels are fine-tuned via the continued proteolytic degradation of VDE (NPQ1) and PsbS (NPQ4), allowing for high electron transfer rate (ETR) and an increased photosynthetic yield. In the absence of the Deg1 protease, the degradation of VDE (NPQ1) and PsbS (NPQ4) is impaired, causing their accumulation and, in turn, an increased level of heat dissipation on account of photochemistry (lower ETR). As a result, the photosynthetic yield is decreased and the development and growth of the plants is compromised.

It is interesting to note that VDE and Deg1 are not only found in the same sub-organellar compartment, and as suggested here, regulate NPQ, but they also share a similar mechanism of activation – through oligomerization that is induced by acidification of the lumen. Although both enzymes have totally different enzymatic activities, they are both monomeric at neutral- and basic pH. Upon acidification of the lumen VDE dimerizes, and the channel connecting the two active sites can accommodate its substrate, violaxanthin, and de-epoxidize it (Arnoux et al., 2009). Acidification of the lumen is critical for activation of Deg1 as well. Protonation of a specific His residue induces a conformational change in the protein that leads to a change in the orientation of its N-terminal α-helix. This, in turn, results in trimerization of the protein and dimerization of trimers, forming the proteolytically-active hexamers (Kley et al., 2011). Thus, both VDE and Deg1 are activated under the environmental conditions that necessitate their activity, to protect and regulate the photosynthetic machinery.

Although VDE and PsbS accumulate to certain levels under most if not all light conditions, we do not know how they are recognized by the constitutively expressed Deg1 protease only when needed. One possibility is that under excess light they themselves get oxidized. This oxidation might induce slight conformational changes that expose termini or loops in the substrates, that are long enough to be recognized by Deg1 (Knopf and Adam, 2018). This and other possibilities will have to be experimentally challenged in future studies.

## METHODS

### Plant material

Four genotypes of *Arabidopsis thaliana* (ecotype Columbia-0) were used in this study: WT, *deg1* single mutant, *deg58* double mutant and *deg158* triple mutant, all of them previously described (Butenko et al., 2018). Seedlings were grown for five weeks under short-day conditions (10 h light / 14 h darkness) at 22/18^0^C (day/night). Photon flux density was ∼75 μmol photons m^-2^ s^−1^, representing normal light (NL) conditions. For high light (HL) treatment, the plants were exposed to ∼750 μmol photons m^-2^ s^-1^ for up to 8 h, starting 1 h after the onset of the light period. For the recovery phase, the HL-treated plants were subjected to additional 24 h under NL. Alternatively, seedlings were grown on half strength Murashige & Skoog agar plates supplemented with 1% sucrose for 15 days at 22^0^C under 100 μmol photons m^-2^ s^−1^, and then transferred to the same liquid medium, with or without 200 μM cycloheximide, for an additional 24-h incubation.

### Chlorophyll fluorescence measurements

All measurements were done on whole plants using the Maxi-Imaging PAM Chlorophyll Fluorometer (Walz, Germany), operated by a dedicated software module (ImagingWin 2.3, Walz). Pulse-modulated chlorophyll *a* fluorescence was recorded as follows: Following dark adaptation for 30 min, the measuring light (9 µmol photons m^-2^ s^-1^) was turned on, and the minimal fluorescence in the dark (*Fo*) was determined. Plants were then exposed to a pulse of saturating light to determine the maximum fluorescence in the dark (*Fm*). After a short delay (40s), made to enable reoxidation following the saturation pulse (SP), the leaves were illuminated with an actinic light (AL) of ∼700 µmol photons m^-2^ s^-1^ on which successive SPs were superimposed at 20-s intervals, enabling determination of the maximum fluorescence in the light-adapted state (*Fm’*). Once steady state has been established, the AL was turned off, the SPs were set at 40-60-s intervals, and the maximal and minimal fluorescence during illumination *(Fm’ and Fo*’) were determined. Rapid light response curves were recorded using a 10-min protocol of successive 20-s cycles of exposure to increasing light intensities, from 1 to 1250 µmol photons m^-2^ s^-1^. All measurements were performed in a dark room at 25^0^C. Parameters calculated included the maximum quantum yield of PSII (*Fv/Fm)*, efficiency of PSII (Y[II]), electron transport rate (ETR) and non-photochemical quenching (NPQ).

### Shotgun proteomic analysis

Frozen leaf samples were crushed in 5% SDS, 0.1M Tris-HCl pH 7.9. The lysates were incubated for 5 min at 96°C, then cleared by centrifugation at 14,000g for 5 min at RT. Proteins were reduced by incubation with 5 mM dithiothreitol (Sigma) for 45 min at 60°C, and alkylated with 10 mM iodoacetamide (Sigma) in the dark for 45 min at RT. The lysates were loaded on a S-trap column and washed with 90% methanol. The proteins were proteolytically digested on the column by incubation with trypsin (Promega) overnight at 37°C, at 50:1 protein/trypsin ratio. Peptides were eluted from the column with 50% acetonitrile and 0.2% formic acid. The samples were vacuum dried and stored in 80°C until further analysis.

Samples were redesolved and loaded in a random order, using split-less nano-Ultra Performance Liquid Chromatography (nanoUPLC; 10 kpsi nanoAcquity; Waters, Milford, MA, USA). The mobile phase consisted of H_2_O + 0.1% formic acid (A) and acetonitrile + 0.1% formic acid (B). Desalting of the samples was performed online, using reversed-phase Symmetry C18 trapping column (180 µm internal diameter, 20 mm length, 5 µm particle size; Waters). Peptides were separated on a T3 HSS nano-column (75 µm internal diameter, 250 mm length, 1.8 µm particle size; Waters) at 0.35 µL/min. They were eluted from the column into the mass spectrometer using the following gradient: 4% to 30%B in 105 min, 30% to 90%B in 5 min, maintained at 90% B for 5 min and then back to 4%B.

The nanoUPLC was coupled online through a nanoESI emitter (10 μm tip; New Objective; Woburn, MA, USA) to a tribrid Orbitrap Fusion Lumos mass spectrometer (Thermo Scientific), using a PicoView nanospray apparatus (New Objective). Data was acquired in data dependent acquisition (DDA) mode, using a top speed method with 3 second cycle time, according to the manufacturer recommendations. MS1 resolution was set to 120,000 (at 200m/z), mass range of 300-1600m/z, Automatic Gain Control (AGC) of 4e5, and the maximum injection time was set to 50msec. Fragmentation was performed in the ion trap, with quadrupole isolation window of 1m/z, AGC of 1e4, dynamic exclusion of 20sec and maximum injection time of 34 msec.

Raw data was processed using MaxQuant version 1.6.0.16. The data were searched with the Andromeda search engine against the TAIR proteome, appended with common lab protein contaminants. Quantification was based on the LFQ method (Cox et al., 2014), based on unique peptides for each protein.

### MS data analysis

Individual intensities were log2 transformed and *Z*-score normalized. Missing values were imputed using normal distribution, with a mean and standard deviation adjusted to resemble low-abundant proteins signals. Proteins were considered for comparative analysis if a protein was identified in at least three out of four replicates. Analysis of Variance (ANOVA), was used to identify significant differences across the biological replicates. The criteria used to denote significantly differentially expressed proteins for further analysis were fold-change >2 and false discovery rate (FDR) correction of 5% (q value < 0.05). The quality of the expressed and differentially expressed data was examined by principal component analysis (PCA). Pairwise Pearson’s correlation was performed to evaluate the dynamic (time-varying) strength of the association between the variables. Unsupervised Hierarchal clustering analysis was performed using Euclidean distance metric and Ward’s Minimum-Variance-linkage agglomerative method. Heat maps were based on K-means clustering, using Pearson correlation coefficient as a distance metric. The optimal number of clusters was computed using the gap statistic method and comprised 1000 Monte Carlo iterations (Tibshirani et al., 2001). Proteins with significantly changed abundances were subjected to gene ontology (GO) annotation using Blast2Go (Conesa and Gotz, 2008) software or the DAVID (Huang da et al., 2009) database. Gene set enrichment analysis (GSEA) of differentially expressed proteins (DEPs) based on Fisher exact test was performed on GO cellular compartments (CC), biological processes (BP) and molecular function (MF). Groups with at least 2 proteins compared to the identified proteins background and a p value <0.05 were considered enriched. For identifications involving uncharacterized/unannotated proteins, sequences were retrieved from the NCBI database and were blasted against Plants/*Arabidopsis thaliana* protein sequences. Analyses were performed using R/Bioconductor (version R.3.4.2), jmp10 software, or Microsoft Excel 2016.

### RT-qPCR analysis

RNA was extracted with the SV Total RNA Isolation System (Promega) then RNA was treated with DNase I (RNase-free) (Ambion, Thermo Fisher Scientific) and purified with PCI (25:24:1) prior to its use in the assays. Reverse transcription was carried out with 200 U of Superscript III reverse transcriptase, in presence of 40 U of RNase OUT (both from Invitrogen, using ∼1 μg of total RNA and 100 ng of a mixture of random hexanucleotides (Promega) and incubated for 50 min at 50 °C. Reactions were stopped by 15 min incubation at 70 °C and the RT samples served directly for real-time PCR. Quantitative PCR (qPCR) reactions were run on a LightCycler 480 (Roche-Diagnostics, Basel, Switzerland), using 2.5 μL of qPCRBIO SyGreen Blue Mix (PCR BIOSYSTEMS, Pennsylvania, USA) and 2.5 μM forward and reverse primers in a final volume of 5 µL. Reactions were performed in triplicate in the following conditions: pre-heating at 95 °C for 10 min, followed by 40 cycles of 10 s at 95 °C, 10 s at 58 °C and 10 s at 72 °C. The nucleus-encoded ubiquitin and tubulin were used as reference genes in the qPCR analyses.

## AUTHOR CONTRIBUTIONS

Z.A. and Z.R. conceived and supervised the project. E.A.-S., L.N. and L.D.S. grew all plants and performed all growth manipulations and treatments. E.A.-S., L.N. and D.C. did all photosynthesis measurements. E.A.-S., L.D.S. and M.K. performed the proteomic and MS data analyses. L.D.S. did RT-qPCR analyses. E.A.-S., L.D.S., D.C., Z.R. and Z.A. analyzed all data. Z.A. wrote the manuscript, with contribution of all the other coauthors.

## SUPPLEMENTAL DATA

The following materials are available in the online version of this article.

**Supplemental Figure 1.** Imaging PAM measurements of PSII maximum quantum yield (*Fv/Fm*).

**Supplemental Figure 2.** Light response curves of photosynthetic electron transport rates (ETRs).

**Supplemental Figure 3.** Pairwise Pearson’s correlation analysis of WT and *deg* mutant plants under NL conditions.

**Supplemental Figure 4.** Principal component analysis (PCA) of the proteomic data of WT and *deg* mutant plants subjected to NL and HL conditions.

**Supplemental Figure 5.** Relative expression of NPQ1 and NPQ4 during high light exposure in WT and *deg* mutants.

**Supplemental Figure 6.** Differential expression of altered proteins in the *deg1* mutant and WT in response to cycloheximide treatment.

**Supplemental Table 1.** Proteomic raw data and analysis of WT and *deg* mutants under NL, HL and recovery.

**Supplemental Table 2.** Proteomic raw data and analysis of WT and *deg1* with or without cycloheximide treatment.

